# Mistimed surveys lead to underestimated migratory bird impacts from wind farms

**DOI:** 10.1101/2025.09.11.675726

**Authors:** Xu Shi, William Holden, Jeremy S. Simmonds, Rohan H. Clarke, Jason W. Chapman, Yang Liu, Richard A. Fuller, Ailsa Kerswell

## Abstract

Accurately estimating the likelihood of occurrence of threatened species is essential for effective impact assessments at development sites. Current survey guidelines often rely on coarse, static species distribution maps, risking misalignment for migratory birds present only seasonally. We evaluated 70 wind farms in eastern Australia to assess the alignment between survey timing and seasonal occurrence of four migratory species. Using eBird-based relative abundance models we classified recommended and actual survey periods as optimal, suboptimal, or poorly timed. Half of all surveys were not optimal, and one fifth missed the species’ potential temporal window of presence altogether. Field detections mostly occurred within modelled optimal or suboptimal windows, validating our approach. Likelihood of occurrence was significantly lower in poorly or suboptimally timed surveys, indicating current guidance for surveying migratory birds at Australian windfarms contains critical shortcomings. Evidence-based alignment with seasonal presence is needed to improve biodiversity assessment guidelines for wind energy development.

## Introduction

The transition to renewable energy is accelerating worldwide, with wind power emerging as an important component of climate change mitigation strategies^1^. Globally, wind energy accounted for 8% of power generation in 2024—a figure projected to triple by 2050^2, 3^. As a cornerstone in global decarbonisation efforts, wind energy is displacing fossil fuel generation and substantially reducing greenhouse gas emissions. Poorly implemented developments, however, can have direct impacts on biodiversity—particularly to wildlife such as birds and bats^4–8^. In the United States, for example, wind turbines are estimated cause mortality of around 200,000 birds annually^9^. Among the most vulnerable are migratory birds^10^, which may collide with turbine blades and towers^11–13^, be forced into energy-intensive detours to avoid wind farms that act as barriers^14, 15^ or experience habitat loss and degradation due to wind farm installations^5, 16^, all of which may exacerbate the ongoing global population declines of migratory birds^9, 17, 18^.

To reduce the risk of approving a wind farm development that could negatively impact migratory birds, robust estimates of the likelihood of occurrence of migratory birds are essential^19^. Accurate assessments can reduce both environmental impacts and the economic costs of curtailment or removal of installed turbines^5^. Globally, environmental impact assessments (EIAs) are widely used to predict potential negative biodiversity outcomes of proposed wind energy projects, often combining desktop analyses with field surveys to produce species-specific evaluations. These pre-construction assessments are critical: they guide formal evaluation of exposure risk to turbine impacts, inform site selection and turbine placement, form the foundation for applying the mitigation hierarchy including avoidance of impacts, and establish a baseline for post-construction monitoring^20^.

However, commonly used assessment tools and datasets often provide information that are spatially coarse and temporally static (e.g., species distribution maps), lacking the seasonal resolution essential for assessing migratory species. This is compounded by the fact that detecting migratory birds in field surveys poses inherent challenges, as many species are only present for brief periods of the year (especially while in transit). Surveys conducted outside these periods risk falsely concluding a species is always absent through the year. Even when species are detected, the misalignment between survey timing and peak periods of abundance may underestimate population size and the associated mortality risk arising from the wind farm should it be built. Consequently, aligning survey timing with peak periods of presence and abundance is essential for producing accurate, ecologically meaningful assessments.

Despite the central role of field surveys in EIAs, such surveys are often constrained by logistical and scheduling factors, including the availability of suitably experienced personnel, unpredictable weather conditions, seasonally variable access to remote sites and commercial deadlines. As a result, many surveys have only narrow temporal coverage. To help address this challenge, some governments have published survey guidelines for migratory species—for example, in Australia via national survey guidelines^21^ and the Species Profile and Threats Database (https://www.environment.gov.au/cgi-bin/sprat/public/sprat.pl). However, these sources lack enforceable requirements or evidence-informed recommendations for optimal survey timing, with suggested survey windows typically spanning the entire period of potential presence across a broad region (e.g. continental or sub-continental scale) that might be wholly inaccurate for a particular location. For example, for the White-throated Needletail *Hirundapus caudacutus*, a long-distance migrant that breeds in Central Siberia and Japan and migrates to Eastern Australia for the non-breeding period, survey guidelines recommend conducting surveys between October and April for Queensland, which spans roughly 2200 km north–south and 1500 km east–west. Yet, the guidelines lack specificity regarding when and where detectability is expected to be highest in any particular part of this large area. This lack of detailed guidance hampers the ability to align fieldwork with the peak occurrence of migratory species, and thus limits the capacity of government and other stakeholders to assess the validity of the impact assessments that are put forward by proponents. Notwithstanding deficiencies in survey effort and accuracy, our knowledge of the migratory dynamics of many species is further limited due to a scarcity of tracking studies and population-level movement data in Australia^22–26^. This limits the reliability of survey guidance that informs impact assessments, and ultimately influences the recovery potential of threatened and migratory species.

Recent advances in data availability and analytical methods offer new opportunities to improve the timing and effectiveness of wildlife surveys. Citizen science platforms such as eBird have transformed avian monitoring by collecting millions of observations contributed by birdwatchers worldwide^27^. Leveraging this data, the eBird Status & Trends project produces weekly, fine-scale model estimates of relative abundance across various species’ ranges^28^. These models incorporate spatial and temporal variation in species occurrence while accounting for observational bias (e.g., variation in observer effort and skill), effectively capturing migratory phenology at high spatial and temporal resolution^29, 30^. As such, they present a valuable tool for benchmarking when and where migratory birds are most likely to occur, providing an empirical basis for defining optimal survey windows.

In this study, we evaluate whether field surveys conducted for onshore wind energy projects are appropriately timed to detect migratory landbird species of conservation concern. We focus on projects in Australia and use four migratory species— White-throated Needletail, Pacific Swift (*Apus pacificus*), Latham’s Snipe (*Gallinago hardwickii*), and Rainbow Bee-eater (*Merops ornatus*) as focal examples. The first three species are long-distance landbird migrants that breed in Asia and spend the non-breeding season (October to April) in Australia each year. Whilst the Pacific Swift and Rainbow Bee-eater are widely distributed across continental Australia, the global populations of both Latham’s Snipe and the White-throated Needletail are nearly wholly constrained to eastern Australian states in the non-breeding season (i.e., the austral summer). The swifts and snipe are also legally protected in Australia as migratory and/or threatened species under national environment law, and therefore must be explicitly assessed in the EIA process for major projects. In addition to their migratory movements, the needletail and swift are obligate aerial insectivores, suggesting they may be particularly vulnerable to wind farm impacts^31, 32^. The Rainbow Bee-eater, in contrast, is a common, widely distributed, and easily detectable medium-distance migrant, endemic as a breeding species to Australia; we included it as a reference species to compare survey effectiveness for a more ubiquitous species with much longer periods of presence.

Specifically, we aim to (i) determine whether field surveys for wind farm EIAs align with species’ periods of peak relative abundance as modelled from eBird data, (ii) assess whether detections, as recorded during field surveys, are more likely when surveys fall within these spatially-explicit optimal survey windows, validating the utility of using eBird data for this purpose, and (iii) discover whether mistimed surveys lead to underestimates of species’ likelihood of occurrence reported in the EIAs. We intend to use these findings to assess the efficacy of current survey guidelines and inform evidence-based practice for wildlife monitoring in EIAs, thereby helping to improve conservation outcomes for migratory birds in Australia.

## Results

A total of 70 planned and constructed wind farms sites were included in this study, ranging from tropical northern Queensland to temperate southern Tasmania (Fig. 1E). eBird models predicted potential presence (max RA > 0.01) at 64 sites for Rainbow Bee-eater, 58 each for White-throated Needletail and Pacific Swift and 30 for Latham’s Snipe (Table S1, Supplementary Material). Impact assessments from these wind farms predicted a known or potential likelihood of occurrence in 47 and 42 sites for White-throated Needletail and Pacific Swift respectively, 37 sites for Rainbow Bee-eater and 17 sites for Latham’s Snipe.

**Figure 1.**
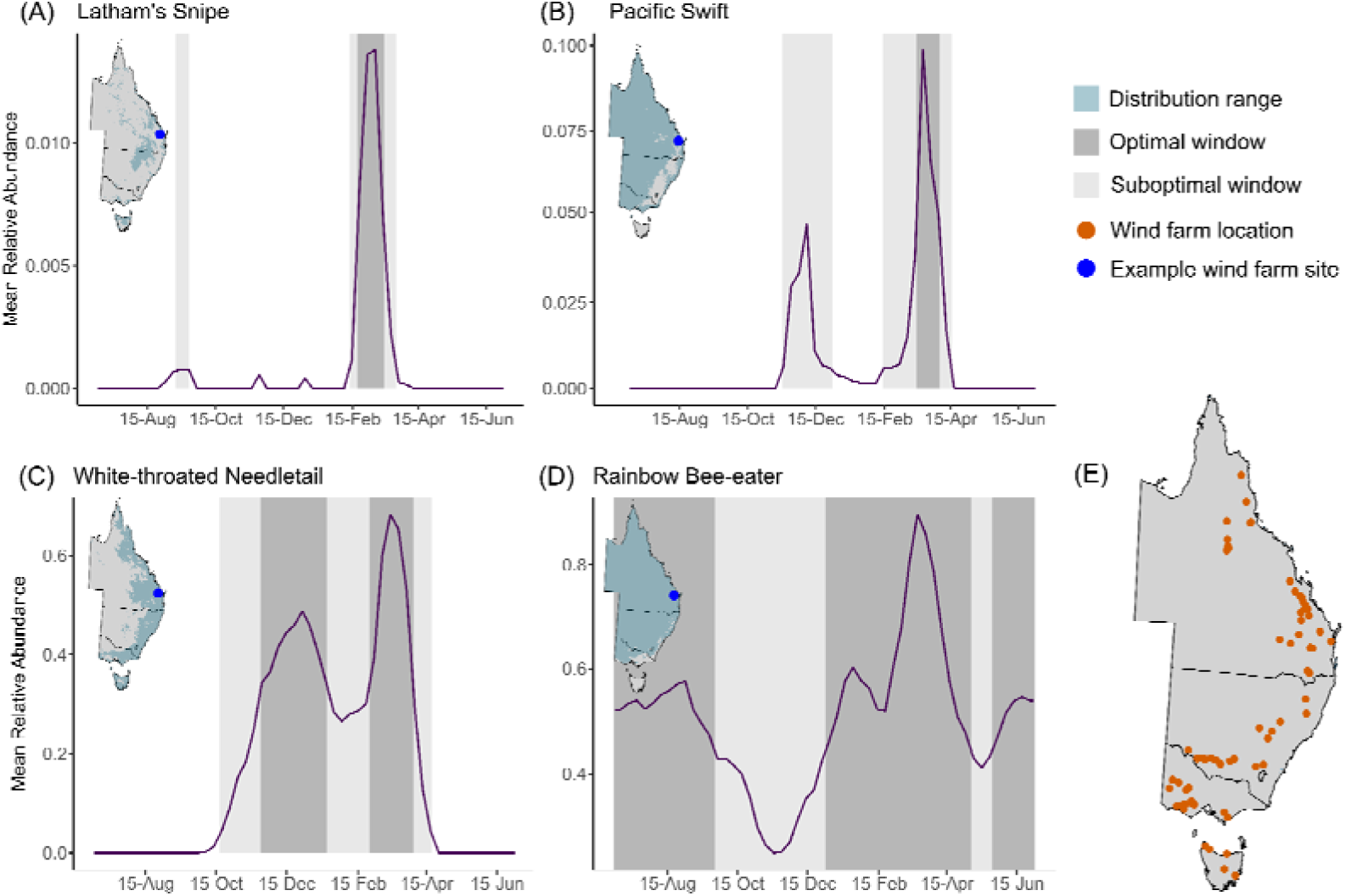
Example of survey windows for the four focal migratory bird species and locations of wind farms in this study. Panels (A–D) show the calculated survey windows at an example wind farm site in southeast Queensland. Optimal survey periods are shown in dark grey, suboptimal periods in light grey, and periods outside these windows would be below optimal survey window (see Methods for definitions). The modelled species range is shown in blue in the inset maps, cropped to eastern Australia. The x-axis spans from July 1 to June 30 to align with the southern hemisphere seasonal cycle and to center peak migratory periods. Note that the example wind farm site (colored in blue) used for window calculation is enlarged for visual clarity in panels (A–D), while (E) displays the spatially accurate 25 km buffer used for analysis with all wind farm sites (colored in orange) used in this study.

Survey window lengths varied markedly among species and along the latitudinal gradient (Fig. 2). The Rainbow Bee-eater had the longest average window if both optimal and suboptimal periods were included (mean = 237 days, SD = 82.8), followed by White-throated Needletail (114 ± 41.1), Latham’s Snipe (106 ± 62.9), and Pacific Swift (84.4 ± 21.9). Rainbow Bee-eater showed the strongest positive association with latitude (slope = 10.1, *p* < 0.001), while Latham’s Snipe showed a strong negative trend (slope = –6.16, *p* < 0.001). Positive relationships were also found for Pacific Swift and White-throated Needletail (slopes = 1.54 and 2.03, respectively; *p* < 0.001). When considering only optimal survey periods, window lengths were substantially shorter for each species (*p* < 0.01, Wilcoxon test): mean window length for Rainbow Bee-eater was 113 ± 65.6 days, while White-throated Needletail, Latham’s Snipe, and Pacific Swift had average optimal windows of 49.2 ± 24.0, 38.5 ± 28.3, and 35.7 ± 12.7 days respectively. Similar latitudinal trends were observed: Latham’s Snipe maintained a strong negative slope (–2.38, *p* < 0.001), and Rainbow Bee-eater and Pacific Swift retained significant positive trends (slopes = 5.53 and 0.42; *p* < 0.001). No significant relationship with latitude was detected for White-throated Needletail (*p* = 0.31).

**Figure 2.**
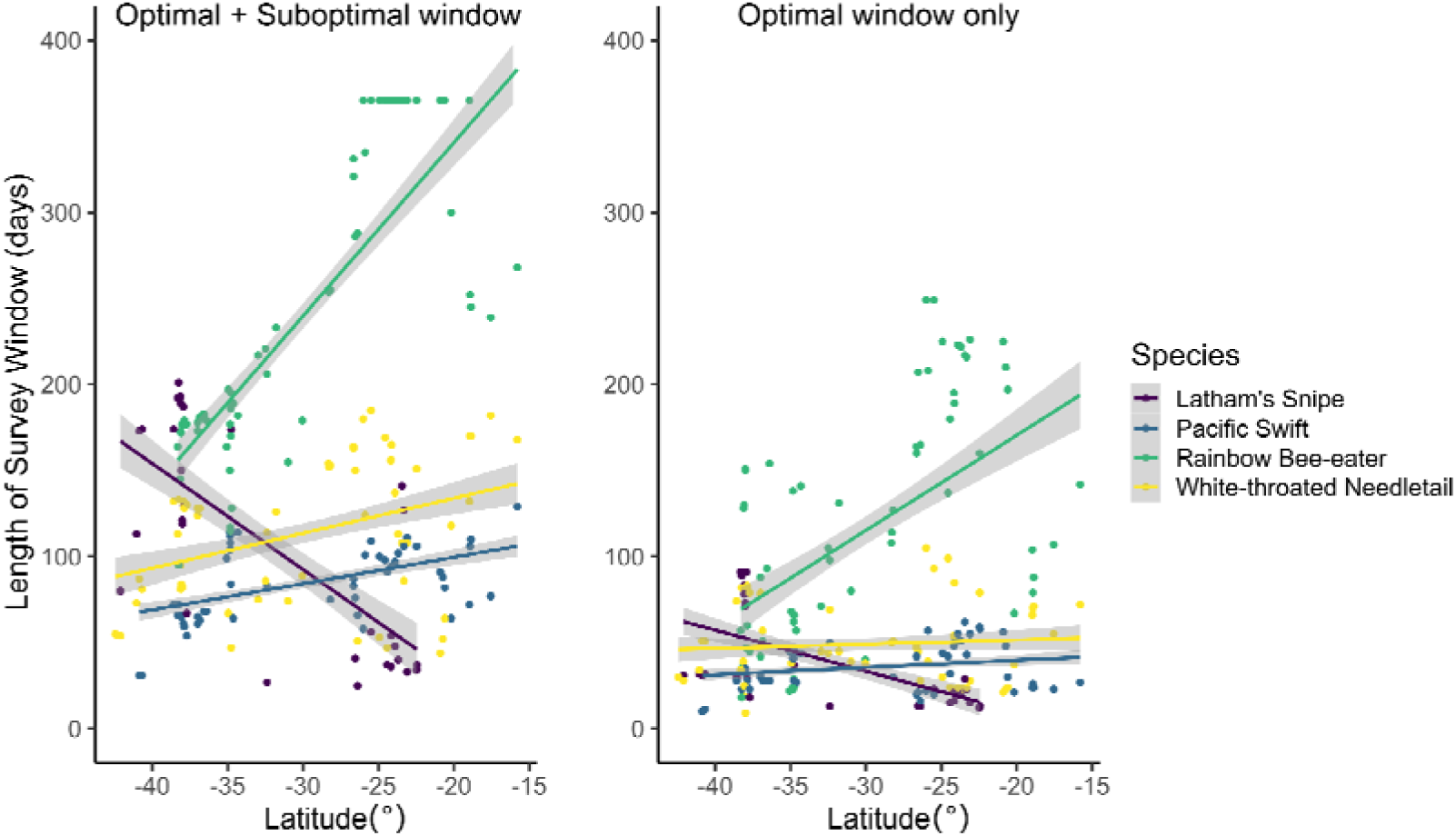
Relationship between survey window length and latitude across wind farm sites. Each point represents the length of the survey window at a given wind farm location at which the respective species was known/potential to occur based on eBird models. The solid line shows the fitted linear regression model, with latitude as the predictor and survey window length as the response variable. Shaded areas indicate the 95% confidence interval of the model fit.

Across all four focal species, 54.2% (114/210) of all surveys were classified as optimal, with suboptimal surveys accounting for 24.7% (52/210) and poorly timed surveys making up 20.9% (44/210) of the total. Survey performance varied among our focal species. The proportion of surveys classified as optimal was highest for Rainbow Bee-eater (72%, 47 out of 64), compared to just 41% for White-throated Needletail (24/58), 47% for Pacific Swift (28/58), and only 45% for Latham’s Snipe (15/33, Fig. 3). Poorly timed surveys were highest for Pacific Swift (31%, 18/58), followed by White-throated Needletail (28%, 14/58) and Latham’s Snipe (18%, 6/33), and lowest for Rainbow Bee-eater (9%, 6/64). The proportion of suboptimal surveys across species spanned a similar range to that of poorly timed surveys, being highest for White-throated Needletails (34%, 24/58) and lowest for Latham’s Snipe (17%, 9/33, Fig. 3).

**Figure 3.**
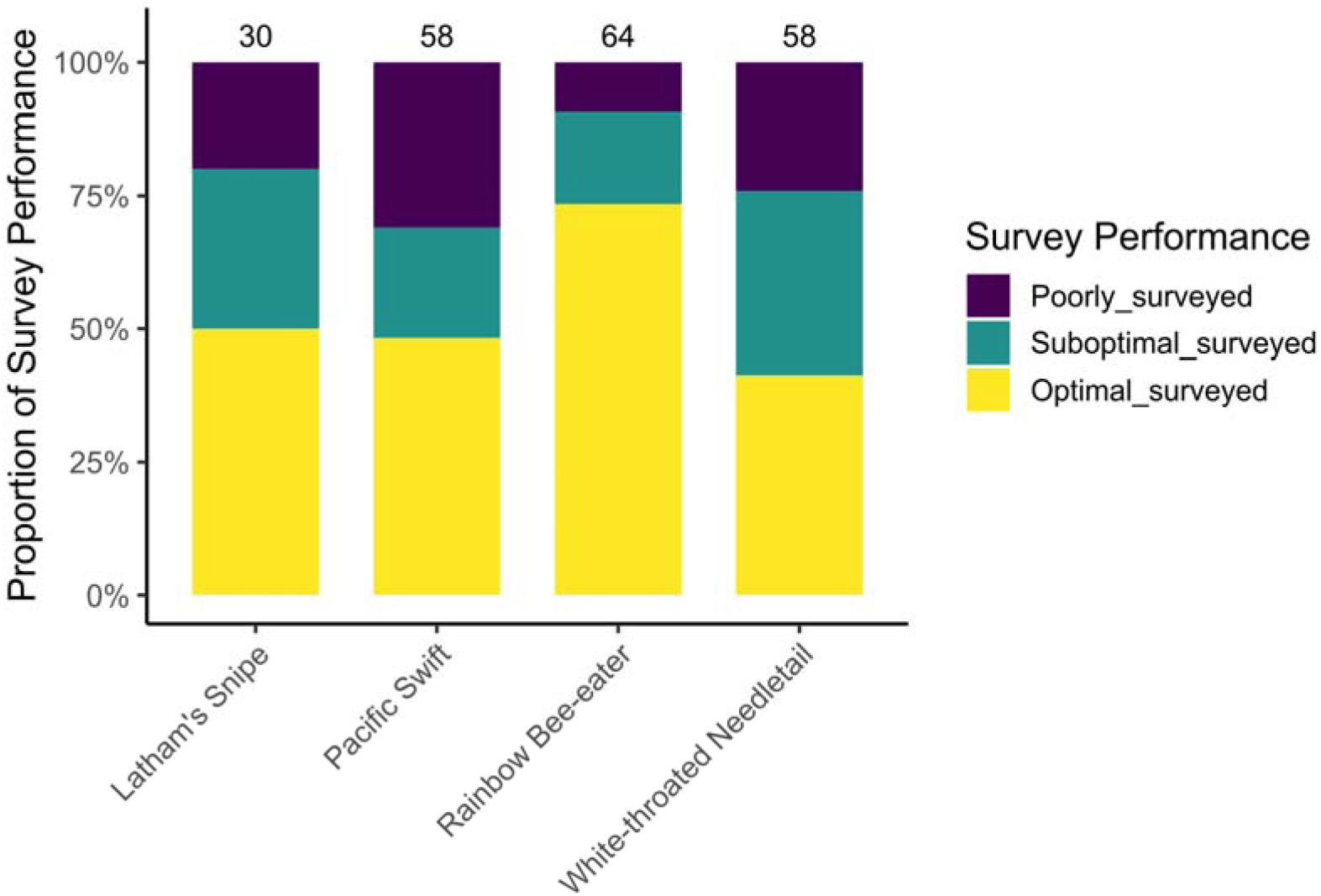
Proportion of types of survey performance for each species. Numbers above columns represent the sample size, defined as the number of wind farms where eBird models predicted potential presence (maximum relative abundance > 0.01).

A total of 30 validation records containing precise observation dates or periods (each may report zero or multiple counts) were available across the four focal species from the environmental assessment reports that we reviewed (Table S1, Supplementary Material). White-throated Needletail had the most validation records (n = 19), followed by Rainbow Bee-eater (n = 10), Pacific Swift (n = 6), and Latham’s Snipe (n = 4). Most species were detected either during optimal survey windows or in suboptimal windows when no optimal survey effort had been conducted (Fig. 4A). Rainbow Bee-eater showed the highest proportion (80%) of detections within these windows, with six records during optimal windows and two during suboptimal windows without optimal effort. White-throated Needletail showed a similar pattern, with nine records in optimal windows and six in suboptimal windows without optimal coverage (15/19, 79%). Three of four detections for Latham’s Snipe and four of six for Pacific Swift occurred in optimal or suboptimal windows without surveys in optimal windows.

**Figure 4.**
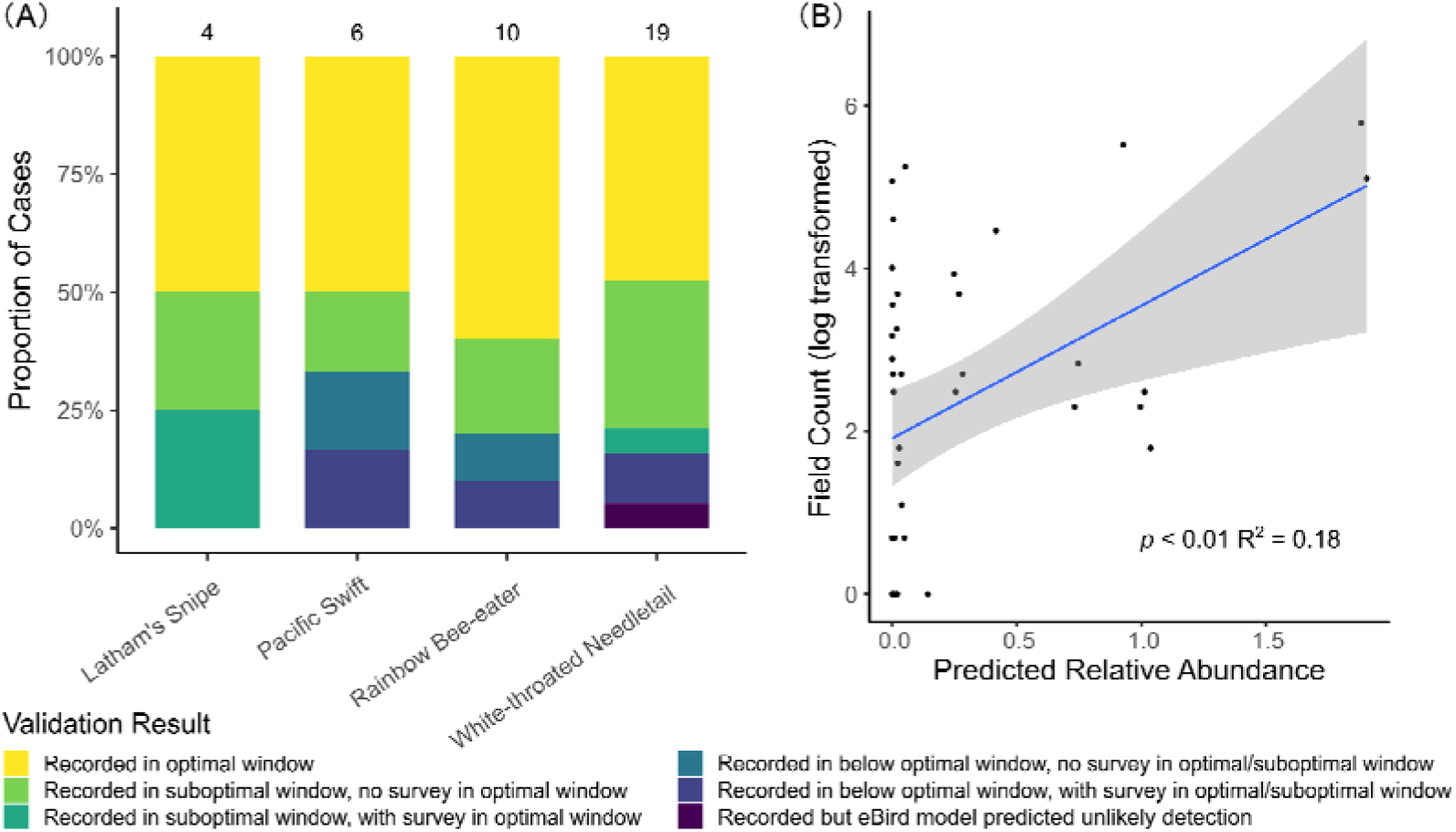
Validation of the survey window classifications using field survey data. (A) Proportions of each validation outcome across species. Numbers above bars indicate the number of wind farm sites with available validation data for each species. (B) Relationship between eBird model-predicted relative abundance and observed field survey counts, pooling together data from all four species (n = 30). The blue line represents the linear regression fit, with the shaded grey area showing the 95% confidence interval. The field counts were log-transformed prior to plotting and modelling.

Detections during periods classified as ‘below-optimal’ were rare, with two instances for each of Rainbow Bee-eater, Pacific Swift, and White-throated Needletail. Additionally, a single White-throated Needletail detection occurred at a time and location where the eBird model predicted a low probability of detection (max RA < 0.01). A total of 43 field counts with associated dates or periods were reported across the four focal species, ranging from one individual (all species) up to a maximum of 327 for a flock of White-throated Needletails (Table S1, Supplementary Material). RA predictions from eBird model for the same day of the year or averaged for the period, and observed counts (log-transformed), showed a significant positive relationship using a linear model (Fig. 4B, *p* = 0.0048, R² = 0.18).

Across all species, records classified with a *Known* likelihood of occurrence (i.e., recorded in field survey) were more prevalent in cases with optimal surveys, especially compared to cases with only poorly surveyed performance (Fig. 5). For instance, 50% (12/24) of optimally surveyed cases for White-throated Needletail were classified as *Known* likelihood of occurrence, compared to 0% (0/14) in poorly surveyed cases and 45% (9/20) in suboptimally surveyed cases. Similarly, for Pacific Swift, *Known* likelihood of occurrence was assigned to 25% (7/28) of optimally surveyed cases, compared to 6% (1/18) in poorly and 17% (2/12) in suboptimally surveyed cases. Latham’s Snipe and Rainbow Bee-eater showed the same pattern: Latham’s Snipe had 33% (5/15) *Known* likelihood of occurrence in optimally surveyed cases, but 0% (0/6) in poorly and 11% (1/9) in suboptimally surveyed cases. Rainbow Bee-eater had the highest proportion, with 62% (29/47) of optimally surveyed cases classified as *Known* likelihood of occurrence, compared to 0% (0/6, note that none of the cases reported likelihood of occurrence) in poorly and 18% (2/11) in suboptimally surveyed cases.

**Figure 5.**
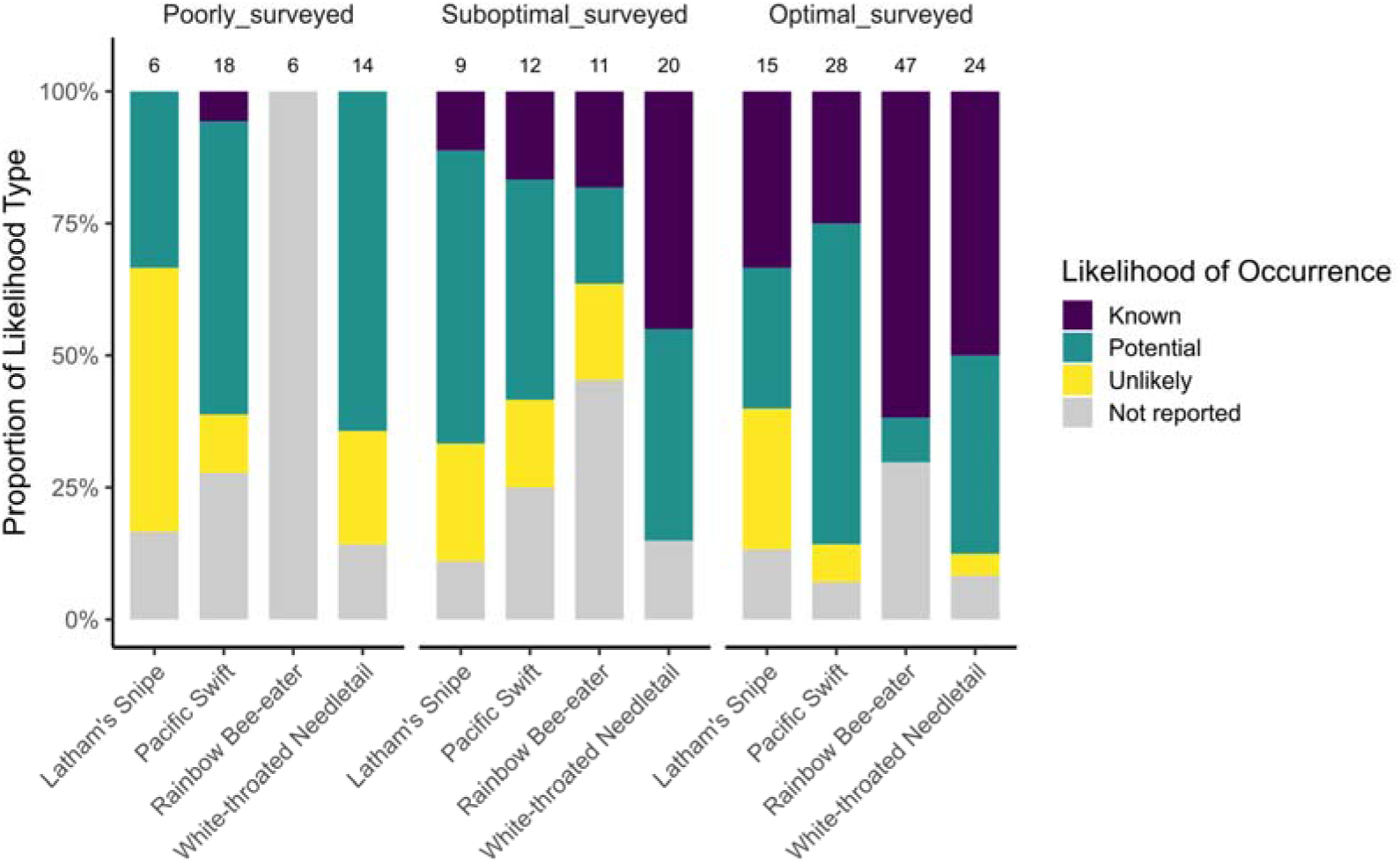
Proportion of likelihood of occurrence categories across survey performance levels for each species. Bars show the relative proportion of wind farms classified as Known, Potential, Unlikely, or Not reported. Likelihood of occurrence categories are extracted from wind farms’ impact assessment reports. Facets represent different survey performance levels. Number on top of each bar represents the total number of cases for each combination of species and survey performance.

Survey timing significantly affected the likelihood of occurrence classification for our focal bird species across wind farms with optimal, suboptimal or poor survey timing producing clear differences (Table 1). Compared to optimally surveyed cases, records from poorly surveyed sites were significantly less likely to receive higher likelihood classifications (Estimate = -1.90, *p* < 0.001). Suboptimal surveys also tended to have lower likelihood classifications, although this effect was marginally significant (Estimate = -0.72, *p* = 0.08). By contrast, latitude, year of submission, and peak relative abundance (RA) predicted by the eBird model were not significant predictors of likelihood of occurrence classification (*p* > 0.2 for all).

**Table 1:** Results from a cumulative link mixed model (CLMM) examining predictors of species likelihood of occurrence across wind farms. The response variable is ordinal with three levels: Unlikely, Potential, and Known. Cases with unreported likelihood were excluded. Predictor variables include survey performance (optimal, suboptimal or poorly surveyed), latitude, year of submission (centered), and the peak relative abundance (RA) predicted by the eBird model throughout the year. Species is included as a random intercept. Estimates are on the log-odds scale.

| Predictor | Estimate | Std. Error | z-value | *p* value |
| --- | --- | --- | --- | --- |
| Survey performance:<br>suboptimal surveyed | -0.72 | 0.42 | -1.73 | 0.08 |
| Survey performance:<br>poorly surveyed | -1.90 | 0.48 | -3.99 | <0.001 |
| Latitude | 0.03 | 0.02 | 1.07 | 0.29 |
| Year of submission | 0.01 | 0.07 | 0.07 | 0.94 |
| Peak RA | 0.68 | 0.53 | 1.27 | 0.20 |

## Discussion

Accurate timing of field surveys is essential for reliably assessing the presence of migratory species in environmental impact assessments. Our analysis of wind farm developments in Australia revealed that almost half of all field surveys conducted for environment impact assessment were suboptimally or poorly timed for long-distance migrants. Of particular concern, a fifth of assessments failed to coincide with the species’ potential presence altogether. Assessments with field surveys outside the optimal seasonal windows were significantly less likely to report a high likelihood of occurrence, indicating that mistimed surveys can lead to a systematic underestimation of species presence and potential risk (e.g. of a project having a significant impact on a species that is neither predicted nor assessed during project planning or EIA). This highlights a critical limitation in current assessment practices and underscores the importance of aligning survey timing with species-specific migratory patterns to fully understand potential impacts and how to avoid or respond to them.

Our findings suggest that current tools for identifying potential species presence and the associated survey guidelines are insufficient to reliably detect migratory birds. The updated national guideline *Onshore wind farm guidance - best practice approaches when seeking approval under Australia’s national environment law*^33^, recommends conducting one bird utilisation survey per season over two years (eight surveys total), aiming to “capture migratory and cryptic bird and bat species”. However, our results demonstrate that the optimal survey window for detecting migratory birds is often narrow in any particular location—typically only 30–40 days. Even if surveyors follow the protocol and conduct one survey every season, there remains a high likelihood that surveys will be mistimed, missing the brief periods when migratory species pass through a site, leading to inefficient use of surveyors and proponents’ effort and resources. This is particularly problematic for highly mobile or flocking species such as Pacific Swifts and White-throated Needletails, for which suboptimal surveys may underestimate peak abundances and poorly timed surveys may fail to detect any presence at all. Consequently, there is a significant risk that, even with careful adherence to the current best-practice guidelines, survey requirements for migratory species are not currently fit for purpose.

Large-scale underestimation of the likelihood of occurrence has implications for the wind energy industry. While final approval decisions consider a range of ecological, social, and economic factors, the assessment of likelihood of occurrence plays a pivotal role in both the initial referral (project risk evaluation) stage of environmental assessments, and EIAs upon which approval decisions are ultimately made. If a species is deemed unlikely to occur at a site, it is not factored into avoidance measures at the design stage, no targeted surveys or species-specific management actions are implemented to mitigate potential impacts, and any residual impacts are not compensated for. As a result, inaccurate classification of a species likelihood of occurrence—particularly at the referral but also during EIA stage—can be problematic. Failing to detect the true presence (i.e., false negatives) for species of conservation concern, can create critical gaps in mitigation and regulatory oversight ^20^. While a single misclassified project may have limited population-level impacts, the cumulative effect of repeated underestimation across multiple developments could directly threaten migratory species populations^34^. Ensuring more accurate and biologically informed assessments from the outset is essential to minimize such oversights and safeguard species^35^. Extending assessments to the more than 300 Australian bird species with population fluctuations indicative of migration^25^ will be a critical step toward addressing knowledge gaps.

We recommend that species-specific survey guidelines be developed for migratory birds, incorporating dynamic survey windows with high spatial and temporal resolution. Our focal species differed markedly in both the timing of peak abundance—which defined optimal survey timing—and survey window duration, with latitude being an important factor. These differences reflect underlying ecological and migratory traits, leading to distinct assessment criteria. For example, Latham’s Snipe, with a non-breeding range centered on south-eastern Australia and Tasmania^36^, had broader optimal survey windows in southern latitudes, but a brief, predictable peak in Queensland aligned with northbound migration in the austral autumn. By contrast, the Rainbow Bee-eater, an Australian breeding endemic and medium-distance migrant, showed shorter optimal windows in the south yet could be surveyed year-round at many Queensland sites. White-throated Needletail and Pacific Swift—both trans-equatorial migrants—had short (<50 days) optimal survey windows across all latitudes, but optimal survey timing shifted progressively from north to south through summer before reversing in autumn, consistent with migratory movements (e.g., Tarburton, 2021). Some similarities in assessment needs between the obligatory aerial foraging swift and needletail suggest species with shared ecological and migratory attributes may be grouped into functional survey groups. Confirmation of such migratory bird groupings has the potential to offer further gains in monitoring effectiveness through the implementation of shared methods.

Our study demonstrates that for migratory species, spatio-temporal models derived from citizen science data can effectively inform survey design. In contrast to Lintott et al. (2016), we did not carry out post-construction field surveys. Instead, we validated our findings using field detections reported in environmental impact assessments. These detections aligned well with the model-defined survey windows and predicted relative abundance, supporting the reliability of our approach. Globally, citizen science observations have been instrumental in identifying and predicting species presence and abundance^37–39^, and increasingly used to understand the ecology of and identify conservation needs for Australian species^40, 41^. Such spatial-temporal models offer a cost-effective and scalable tool for improving assessments of migratory species presence and abundance. In Australia, where tracking data and detailed knowledge of migratory routes and timing are still lacking for many species, the ‘wisdom of the crowd’ offered by large-scale citizen science initiatives becomes especially valuable^25, 42^. We suggest that such resources be integrated into revised survey guidelines in Australia to better support ecologists and wind energy developers in identifying periods of likely occurrence and peak abundance of migratory birds at project sites. Incorporating these data-driven insights would enhance the accuracy of impact assessments and contribute to more informed and responsible development planning.

While our approach offers a systematic framework for evaluating survey timing of migratory species, some limitations persist. Spatial gaps are common in eBird models, particularly for cryptic or hard-to-detect species and in less-populated regions such as inland Australia^43^. Additionally, our analysis focused solely on the temporal alignment of surveys and did not account for important variables such as survey method, observer effort (e.g., how often the sky was scanned for swifts and needletails), or the experience level of personnel, all of which may significantly influence detectability, but can be difficult to quantify and are often unreported in impact assessments. Additionally, we found that smaller buffers resulted in narrower, more variable windows, while larger buffers smoothed seasonal curves and expanded window length. Therefore, our definition of survey windows, and subsequent evaluation of survey performance and likelihood of occurrence, are sensitive to buffer size. Nevertheless, about three-quarters of all migratory species records detected during field surveys fell within the optimal window, or within suboptimal windows when no surveys in the optimal window was conducted, suggesting that the 25 km buffer used in this study is reasonable, and can be used as a starting point for future application of our approach before fine-tuning for a species- or project-specific buffer range.

Although this study focuses on wind farm projects in Australia, the methodology can be used to guide field surveys and impact assessments for migratory birds for a wide range of development projects (e.g., buildings, power lines, solar farms) in any region globally with sufficient eBird model output. As temporal and spatial eBird data coverage continues to grow, so will our understanding of spatiotemporal movement patterns that underpins guidance for EIAs. Our study specifically examines medium to long-distance migrants, while many Australian species are partial migrants that undertake shorter intra-continental movements, with often complex movement patterns that are poorly understood^42, 44, 45^. Survey windows may differ subtly yet critically between regions with resident and migratory populations, and our approach can help optimize surveys for these species. Currently, our method evaluates the presence of species at single wind farm sites. However, as wind farm development grows rapidly—not only in Australia but also in regions like East Asia—species such as Pacific Swifts and White-throated Needletails will encounter an increasing number of wind farms along their migration routes^46^. Future work to assess cumulative and population-level impacts would be instrumental in guiding effective conservation strategies in the face of expanding renewable energy infrastructure.

## Methods

### Environmental impact assessment process in Australia

In Australia, the national-level regulatory framework of industrial development is the Australian *Environment Protection and Biodiversity Conservation Act 1999*^47^. Wind energy proponents are required to submit a referral before wind farm construction can be approved, if the project has, will have, or is likely to have a significant impact on Matters of National Environmental Significance (MNES). The referral includes an evaluation of the proposed project’s potential impact on MNES, including migratory species, threatened species, and ecological communities that have been listed under legislation. Any proposed development deemed likely to have a ‘significant impact’ on MNES must be referred to the Department of Climate Change, Energy, the Environment and Water (DCCEEW)—the Australian Government environmental regulator—for detailed environmental impact assessment, ultimately leading to the project’s approval or rejection.

To identify MNES potentially occurring in project areas, two main methods are used: review of desktop data sources and information, including the Australian Government’s Protected Matters Search Tool, and field surveys. The search tool, provided by DCCEEW (www.dcceew.gov.au/environment/epbc/protected-matters-search-tool), generates desktop reports of the species both present and potentially present in a user-defined (i.e., assessment) area. Species with recent records inside the user-defined area are listed as ‘known to occur’, while those within range or suitable habitat but lacking recent records are listed as ‘may occur’. This desktop assessment informs the need for targeted field surveys. Field surveys, typically arranged by project proponents and conducted by contracted environmental consultants, validate these desktop predictions of species presence, as well as determine abundance of focal species, and habitat amount, condition, and configuration in and near the assessment area. The combined results from desktop assessment and field surveys are used to generate a likelihood of occurrence where a species will be deemed ‘known to occur’ if recorded during field surveys, and ‘potential to occur’ or ‘unlikely to occur’ if not recorded during field surveys depending on whether suitable habitat and historical records exist. These evaluations ultimately contribute to an assessment of the significance of the potential impact of the project on population and habitat viability for listed species.

### Pre-construction survey data collection

We accessed wind farm project referrals and associated environmental impact assessment documents from the EPBC Act Public Portal (https://epbcpublicportal.environment.gov.au/all-referrals/). Wind farm projects were identified by filtering for ‘Energy Generation and Supply (Renewable)’ under ‘Industry Type’ and searching keywords such as ‘wind’, ‘wind energy,’ and ‘wind farm’. We constrained our study to the eastern half of Australia, namely Tasmania, Victoria, Australian Capital Territory, New South Wales, and Queensland, where most existing and planned wind farms are located. For each project, we recorded the EPBC project number, the location, the date of project submission to the EPBC portal, project status at the start of our analysis, the state in which the project is located, as well as the Protected Matters Search Tool results for the four species and the proponent’s evaluation of the significance of impact on these species if relevant. From the associated environmental assessment reports or ‘Bird and Bat Utilisation Survey Reports’, we extracted the start and end dates of field surveys. We included all surveys that potentially recorded birds, including bird utilisation surveys, general fauna baseline surveys and flora and fauna monitoring surveys.

Where applicable, we also recorded whether any of the focal species were observed during the surveys and, if reported, the date and number of individuals detected. When multiple detections occurred on the same day, only the highest count was retained to avoid double-counting. To ensure methodological consistency and deliver a consistent, objective and proportional evaluation of proponent submissions in keeping with best-practice guidelines for regulatory assessments in the impact assessments, we limited our dataset to projects submitted after the publication of the *Referral guideline for 14 birds listed as migratory species under the EPBC Act* in 2015^21^ and up to November 2024 when the present study began. Projects with vague survey timing (e.g., only mentioning ‘summer’ or a specific month) were excluded, as were those with a status of ‘project withdrawn’ or ‘clearly unacceptable.’ After these filters, a total of 70 wind farm projects remained in the dataset. A full list of wind farms and their associated survey timing and results is provided in the appendix.

### Extract optimal survey timing and compare with pre-construction surveys

We obtained weekly relative abundance data as well as static range maps from the eBird Status & Trends project for the four species as raster files with a 3 × 3 km spatial resolution using the R package ‘ebirdst’ (version 2.2021.3). We used a spatial buffer centered around each wind farm location to calculate the mean relative abundance trend from the gridded eBird data. Typical wind farm sizes (e.g., 10 - 50 km^2^) are too small to reliably intersect enough pixels given the resolution of eBird data. Small buffers (e.g., 5 km radius around the wind farm location) may better reflect local trends but risk data gaps and underrepresenting habitat variation, while large buffers (e.g., 50 km radius) improve data availability but dilute localized phenology. To determine a suitable buffer size, we examined how optimal survey window length varied with buffer size. Optimal survey window length refers to the number of days within a year classified as an optimal period for detecting a given species, based on seasonal patterns of occurrence (see next paragraph for details). We calculated the relationship between buffer size and mean optimal window length for all wind farms across a range of buffer radius (0 to 100 km, Figure S1, Supplementary Material). For all species, window length increased with buffer size, though with diminishing returns. The rate of increase slowed notably for Latham’s Snipe and Pacific Swift, but less so for Rainbow Bee-eater and White-throated Needletail. To identify an inflection point—beyond which further increases in buffer size yielded little additional gain—we fitted a Generalized Additive Model (GAM) and calculated the first local minimum in its first derivative. This point occurred at approximately 17 km for Latham’s Snipe and 31 km for Pacific Swift. To balance spatial precision with consistency across species, we selected a 25 km buffer for all subsequent analyses.

For each 25 km-buffered wind farm location, we extracted time series of relative abundance from the weekly eBird spatial models for each species and averaged the values within each buffer to produce a mean relative abundance. The weekly data were then linearly interpolated to a daily (day-of-year) resolution to align with survey periods from the environmental assessment reports, which are typically reported using start and end dates. From each daily relative abundance time series, we identified the peak value and calculated two thresholds: 50% and 5% of the peak. Firstly, if the number of peak relative abundance was less than 0.01 (in unit of number of individual birds expected to be observed on a standard checklist), we classified the species as ‘unlikely to occur’ at that location, otherwise as ‘potential to occur’. For those that were potential to occur, we defined the period during which mean relative abundance falls between 50% and 100% of the peak as the ‘optimal survey window’, representing the time when the species is most abundant. The period between 5% and 50% of the peak was classified as a ‘suboptimal survey window’, indicating the species is present but in lower numbers. When relative abundance dropped below 5% of the peak, we considered this ‘below optimal survey window’, suggesting the species is unlikely to be present. We reported the length of each species’ optimal and suboptimal survey window in relation to latitude.

We subsequently compared the survey dates for each wind farm to the survey windows defined above. First, for each wind farm’s field survey efforts, we compiled survey dates from different years into a list of unique days of the year (DOY) when surveys were conducted. We then compared this list to the optimal and suboptimal survey windows. If at least one surveyed DOY fell within the optimal window, the site was classified as ‘optimally surveyed’. If no surveyed DOYs fell within the optimal window but at least one fell within the suboptimal window, the site was classified as ‘sub-optimally surveyed’. If none of the surveyed DOYs overlapped with either the optimal or suboptimal windows, the site was classified as ‘poorly surveyed’.

We also validated our classification of survey windows using actual species detections from field surveys. For wind farms where a species was observed, we compiled the specific dates or periods of these observations and compared them to our defined survey windows to assess whether detections aligned with periods of higher predicted relative abundance. Based on this comparison, each observation was categorized into one of several outcomes: If the species was observed on dates within the optimal survey window, it was classified as ‘recorded in optimal survey’. If the species was observed during the suboptimal window and no surveys were conducted during the optimal window, it was classified as ‘recorded in suboptimal survey, no optimal survey conducted’. In cases where the species was observed during the suboptimal window despite surveys also having been conducted during the optimal window, it was categorized as ‘recorded in suboptimal survey’. Similarly, if the species was observed during the below optimal window and no surveys occurred during the optimal window, the case was classified as ‘recorded in below optimal survey, no optimal survey conducted’. If the species was observed during the below optimal window while surveys were also conducted in the optimal period, it was labeled ‘recorded in below optimal survey’. Finally, if the species was observed at a site where our model predicted it was unlikely to occur (peak RA < 0.01), the observation was categorized as ‘seen but eBird model predicted unlikely detection’. A detailed description of each of the classifications is provided in Table S2 of the Supplementary Material.

We also examined the relationship between the number of individual birds observed on a given date and the predicted relative abundance for the same day of the year using a linear regression model. All available observations were included in the analysis, meaning that multiple observations of a species from different dates within the same field survey were treated as separate data points. In cases where observations were reported over a period rather than a specific date, we used the average relative abundance across that period for comparison. We did not differentiate between species in this analysis, as the relative abundance estimates are standardized and thus comparable across species. We applied a logarithmic transformation to the count data to reduce the influence of high counts before regression with relative abundance.

To investigate whether variation in survey performance influenced the assessment of likelihood of occurrence at wind farm sites, we first compared the proportion of each type of likelihood of occurrence across survey performance levels. We then fitted a cumulative link mixed model (CLMM) using the ‘ordinal’ package in R. The response variable was the likelihood of occurrence, an ordinal variable with three ordered levels: Unlikely, Potential, and Known. We used the following predictor variables: survey performance, which is a categorical variable that included optimal, suboptimal, and poorly surveyed; latitude of the wind farm; year of project submission (centered around the mean); and peak relative abundance (RA) from the eBird model at the site throughout the year. Species was included as a random intercept to account for species-level variation in reporting patterns. We used a logit link function for the model, and all estimates are reported on the log-odds scale, where positive coefficients indicate an increased likelihood of higher occurrence categories. Wind farm cases with missing likelihood of occurrence information (‘Not reported’) were excluded from the model.

## Data Availability

The migratory bird relative abundance data used in this study are derived from the eBird Status and Trends products, available at https://ebird.org/science/status-and-trends.

## Notes

### Competing Interest Statement

The authors have declared no competing interest.

