## Supplementary Material for "Mistimed surveys lead to underestimated migratory bird impacts from wind farms"

**Supplementary Information**

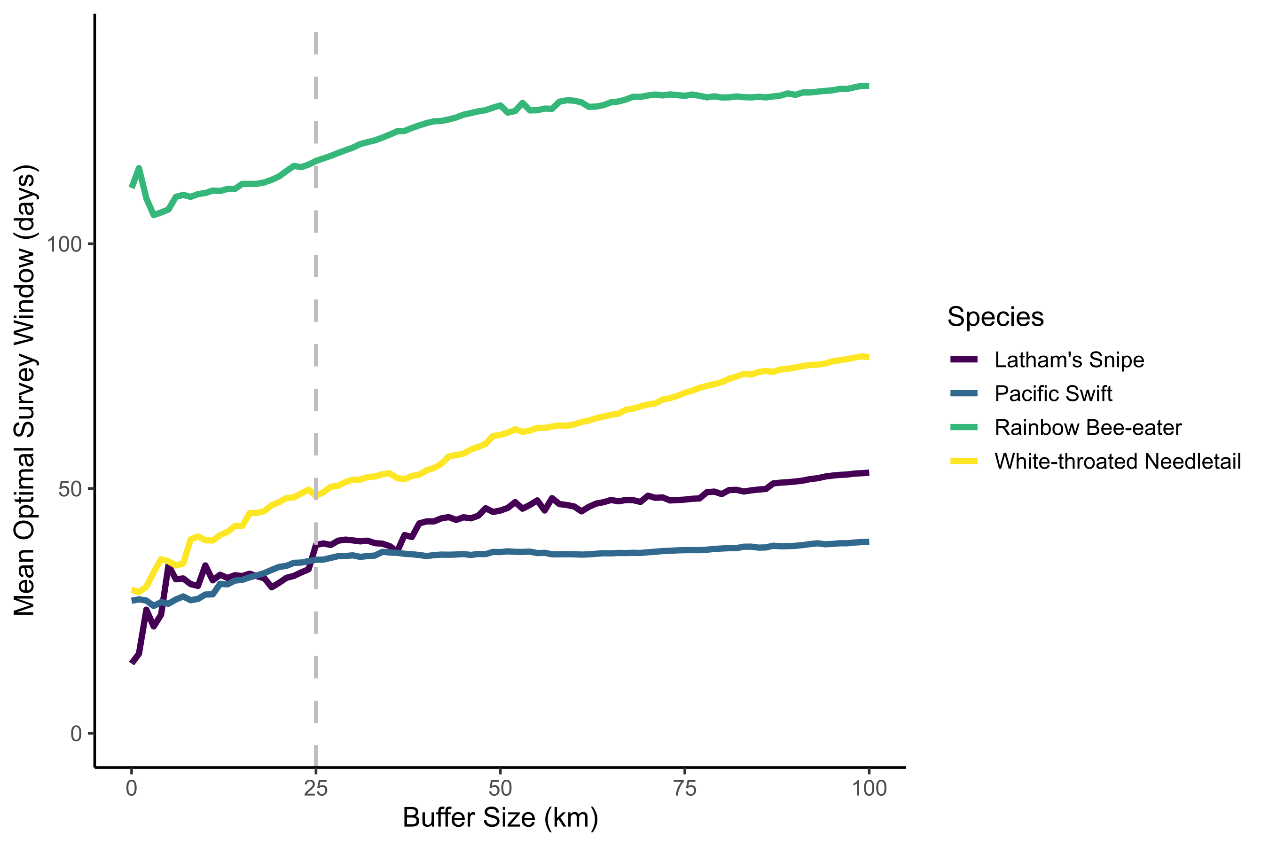

Figure S1. Relationship between buffer size around the wind farm location and the mean optimal survey windows across all wind farm locations. Grey vertical line indicates the 25 km buffer used in this study.

Table S1. Summary of species occurrence across data sources at wind farm sites. “PMST” shows the number of wind farms where species were classified as known or may occur by the Protected Matters Search Tool. “eBird Model Prediction” indicates sites where eBird models predicted potential presence (max RA > 0.01). “Recorded during survey” refers to wind farms with detections in survey reports. “Number of counts with dates reported” is the total number of individual detections with counts and dates (one site may report multiple counts).

| Species | PMST | eBird Model Prediction | Recorded during survey | Number of counts with dates reported | Flock size range |
| --- | --- | --- | --- | --- | --- |
| Latham’s Snipe | 65 | 30 | 7 | 4 | 1-2 |
| Pacific Swift | 62 | 58 | 10 | 6 | 1-51 |
| Rainbow Bee-eater | 47 | 64 | 37 | 3 | 1-12 |
| White-throated Needletail | 58 | 58 | 24 | 30 | 1-327 |

Table S2. Description of survey outcome classifications used in this study.

| Timing of actual observation | Survey conducted in optimal window | Description | Classification |
| --- | --- | --- | --- |
| Within Optimal Survey Window | Yes or No | Species observed during the optimal survey period. | Recorded in optimal survey |
| Within Suboptimal Window | No | Species observed during suboptimal period; no optimal surveys conducted. | Recorded in suboptimal survey, no optimal survey conducted |
| Within Suboptimal Window | Yes | Species observed during suboptimal period despite optimal surveys also conducted. | Recorded in suboptimal survey |
| Within Below Optimal Window | No | Species observed during below optimal period; no optimal surveys conducted. | Recorded in below optimal survey, no optimal survey conducted |
| Within Below Optimal Window | Yes | Species observed during below optimal period despite optimal surveys also conducted. | Recorded in below optimal survey |
| Any time (any window) | N/A | Species observed where modelled peak relative abundance (RA) < 0.01 (unlikely occurrence). | Seen but eBird model predicted unlikely detection |
